# Genetic analysis of the novel SARS-CoV-2 host receptor *TMPRSS2* in different populations

**DOI:** 10.1101/2020.04.23.057190

**Authors:** Roberta Russo, Immacolata Andolfo, Vito Alessandro Lasorsa, Achille Iolascon, Mario Capasso

## Abstract

The infection coronavirus disease 2019 (COVID-19) is caused by a virus classified as severe acute respiratory syndrome coronavirus 2 (SARS-CoV-2). At cellular level, virus infection initiates with binding of viral particles to the host surface cellular receptor angiotensin converting enzyme 2 (ACE2). SARS-CoV-2 engages ACE2 as the entry receptor and employs the cellular serine protease 2 (TMPRSS2) for S protein priming. TMPRSS2 activity is essential for viral spread and pathogenesis in the infected host. Understanding how TMPRSS2 protein expression in the lung varies in the population could reveal important insights into differential susceptibility to influenza and coronavirus infections. Here, we systematically analyzed coding-region variants in *TMPRSS2* and the eQTL variants, which may affect the gene expression, to compare the genomic characteristics of *TMPRSS2* among different populations. Our findings suggest that the lung-specific eQTL variants may confer different susceptibility or response to SARS-CoV-2 infection from different populations under the similar conditions. In particular, we found that the eQTL variant rs35074065 is associated with high expression of *TMPRSS2* but with a low expression of the interferon (IFN)-α/β-inducible gene, *MX1*, splicing isoform. Thus, these subjects could account for a more susceptibility either to viral infection or to a decrease in cellular antiviral response.

## Introduction

In December 2019 a new infectious respiratory disease emerged in Wuhan, Hubei province, China.^1–3^ Subsequently, it diffused worldwide and became a pandemic. The World Health Organization (WHO) has officially named the infection coronavirus disease 2019 (COVID- 19), and the virus has been classified as severe acute respiratory syndrome coronavirus 2 (SARS-CoV-2). The mechanism of infection of SARS-CoV-2 is not yet well known; it appears to have affinity for cells located in the lower airways, where it replicates.^4^ COVID- 19 cause a severe clinical picture in humans, ranging from mild malaise to death by sepsis/acute respiratory distress syndrome.

At cellular level, virus infections initiate with binding of viral particles to host surface cellular receptors.^5,6^ Receptor recognition is therefore an important determinant of the cell and tissue tropism of a virus. Recently, human angiotensin converting enzyme 2 (ACE2) was reported as an entry receptor for SARS-CoV-2.^3^ Moreover, the spike (S) protein of coronaviruses facilitates viral entry into target cells. Entry depends on binding of the surface unit, S1, of the S protein to a cellular receptor, which facilitates viral attachment to the surface of target cells. SARS-CoV-2 engages ACE2 as the entry receptor and employs the cellular serine protease 2 (TMPRSS2) for S protein priming.^5^ TMPRSS2 activity is essential for viral spread and pathogenesis in the infected host.^7–10^ TMPRSS2 as a host cell factor is critical for spread of several clinically relevant viruses, including influenza A viruses and coronaviruses.^7,10,11–16^ TMPRSS2 is a cell surface protein that is expressed by epithelial cells of specific tissues including those in the aerodigestive tract. It is dispensable for development and homeostasis and thus, constitutes an attractive drug target.^13^ In this context, it is noteworthy that the serine protease inhibitor camostat mesylate, which blocks TMPRSS2 activity has been approved in Japan for human use, but for an unrelated indication.^10,14^

Due to the crucial role of TMPRSS2 in the viral infection, we analyzed its genetic landscape in different populations trying to find a possible genetic predisposition to SARS-CoV-2 infection.

## Results

To systematically investigate the candidate functional variants in *TMPRSS2* and the allele frequency (AF) differences between 17 populations with different ethnic origin, we analyzed all the 1025 variants in *TMPRSS2* gene region downloaded from the gnomAD browser and annotated with 34 pathogenic variant scores (Supplementary Table S1). The locus region comprises 496 non-coding and 520 coding variants. The AFs of all the variants located in the coding region of *TMPRSS2* in different largescale genome databases were summarized in Supplementary Table S2. Forty-three loss-of-function (LoF) variants were annotated in gnomAD in the *TMPRSS2* gene locus. The benign variants were classified by using a combination of the three algorithms VEST3, REVEL, and RadialSVM and the pathogenic ones by other three algorithms MutationTaster, Mcap, and CADD as recently suggested.^15^ The 26% (88/334) of non-synonymous variants has been classified as pathogenic. All these variants are located along the entire coding region of the gene (Figure 1a), and both missense pathogenic and LoF variants exhibit very low overall AFs (Figure 1b). This finding agrees with the recommended benign frequency cut-off of 0.0001 for *TMPRSS2* gene, as derived from the Varsome database (https://varsome.com/). Nevertheless, the distribution of AFs across the different populations showed the highest percentage in African (AFR) population within LoF variant class. Similarly, Swedish population exhibited the highest AF for the LoF variants among the Europeans (Figure 1c). When we looked at non-synonymous pathogenic variants, we observed the highest AF among Ashkenazi Jewish (ASJ) individuals (Figure 1b), while Finnish (FIN) showed the highest AF among European subpopulations (Figure 1c). The AFs of non-synonymous variants classified as benign (198/334, 59.3%) were similarly distributed among the different populations (Figure 1b-c). The average AFs for all types of variants here investigated are summarized in Supplementary Table S3.

**Figure 1.**
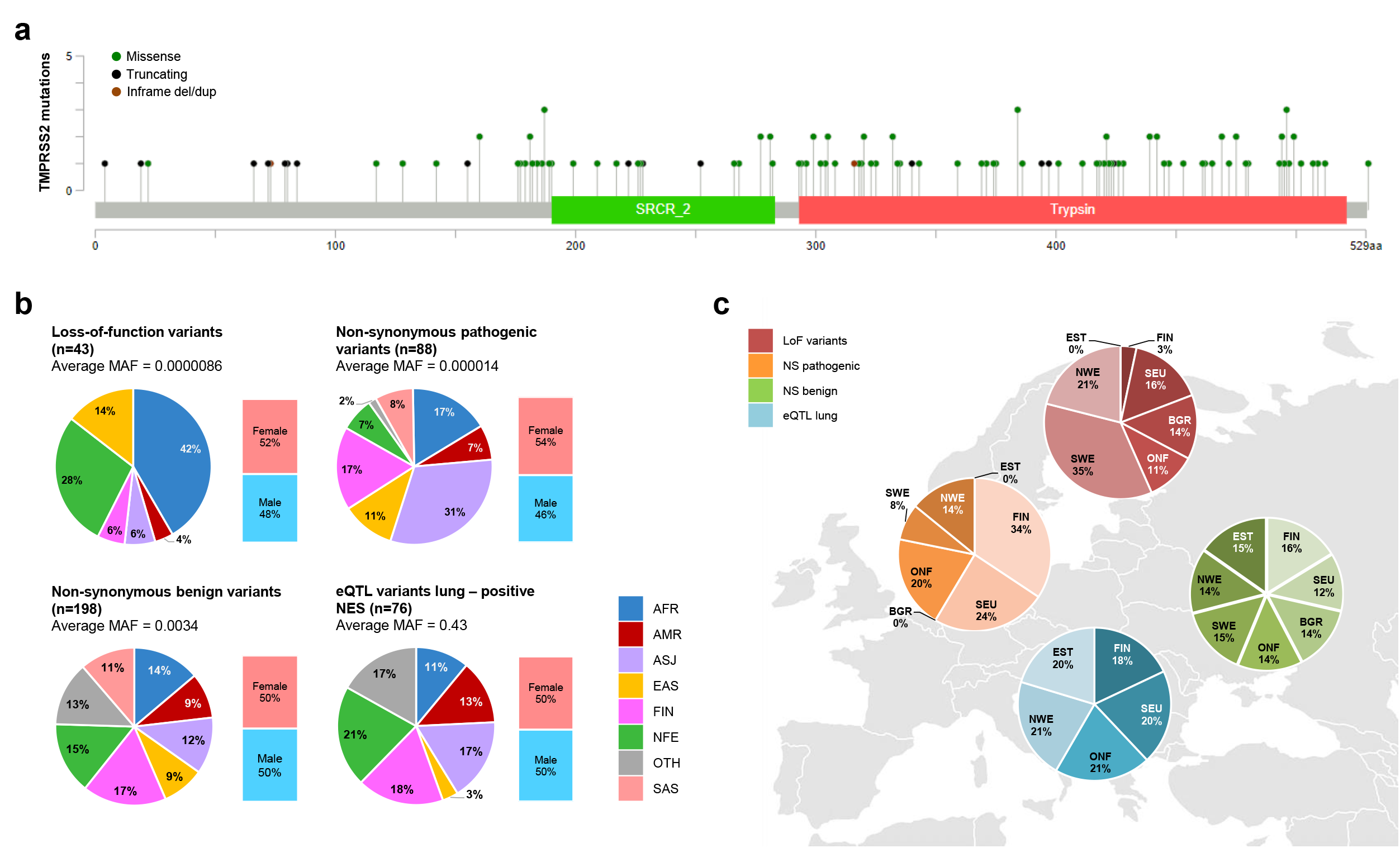
Analysis of the coding-region variants and the eQTL variants for *TMPRSS2* locus in different populations. **a)** Lollipop diagram of TMPRSS2 protein structure and schematics of 88 pathogenic and 33 loss-of-function variants identified in gnomAD database. Mutation diagram circles are colored with respect to the corresponding mutation types. In case of different mutation types at a single position, color of the circle is determined with respect to the most frequent mutation type. Mutation types and corresponding color codes are as follows: missense variants are in green, truncating variants are in black (https://www.cbioportal.org/mutation_mapper). **b)** The allele frequency distribution of non-synonymous pathogenic, benign, loss-of-function, and eQTL-lung variants (positive NES) of *TMPRSS2* in different populations. The colors of each pie chart indicate different populations as shown in the color code legend. AFR, African/African American; AMR, Latino/Admixed American; ASJ, Ashkenazi Jewish; EAS, East Asian; FIN, Finnish; NFE, Non-Finnish European; SAS, South Asian; OTH, Other (population not assigned). The histograms show the AF distribution in the overall gnomAD population stratified according to the gender. **c)** The allele frequency distribution of non-synonymous pathogenic, benign, loss-of-function, and eQTL-lung variants (positive NES) of *TMPRSS2* in different European populations. The colors of each pie chart indicate different types of variants as shown in the color code legend. FIN, Finnish; SEU, Southern European; BGR, Bulgarian; ONF, Other non-Finnish European; SWE, Swedish; NEW, North-Western European; EST, Estonian. The histograms show the AF distribution in the overall gnomAD population stratified according to the gender.

We also investigated, throughout the GTEx database, the distribution of the expression quantitative trait loci (eQTL) for *TMPRSS2* (Supplementary Table S4). Indeed, we found 203 unique and significant (FDR<0.05) eQTL variants for *TMPRSS2* in five different tissues divided as follows: 136 (66.9%) in lung, 56 (27.6%) in testis, 9 (4.4%) in prostate, 1 (0.5%) in ovarian and in thyroid (0.5%) tissue.

*TMPRSS2* is highly expressed in prostate, testis, stomach, colon-transverse, pancreas, and in tissues of the respiratory tract, as bronchus, pharyngeal mucosa, and lung. However, no difference in gene expression between male and female was observed for non-gender specific tissues (Supplementary Figure S1). The AFs of the 136 eQTL-lung variants were compared among different populations, but no substantial differences in AF distribution was observed (Supplementary Figure S2). Nevertheless, the average AF of 76 eQTL-lung variants with positive normalized effect size (NES) was higher in European populations (FIN, 0.463; NFE, 0.541), whereas the average AF of these variants in East Asian (EAS) population was much lower (0.085) (Figure 1b and Supplementary Table S3).

Interestingly, the top 25 variants (NES > 0.1) were in a genomic region that includes both *TMPRSS2* and *MX1* genes. In particular, the most significant eQTL variant rs35074065 is located in the intergenic region between the two genes (distance = 2379 from *MX1*; distance = 2958 from *TMPRSS2)* (Figure 2a) and shows the lowest AF in EAS (0.0049) population (Figure 2b). Of note, this variant is associated with high expression of both *TMPRSS2* and *MX1* in lung tissue (Figure 2c). Notably, the same variant is also a splicing (s)QTL associated with low expression of *MX1* splicing isoform in different tissues (Figure 2d).

**Figure 2.**
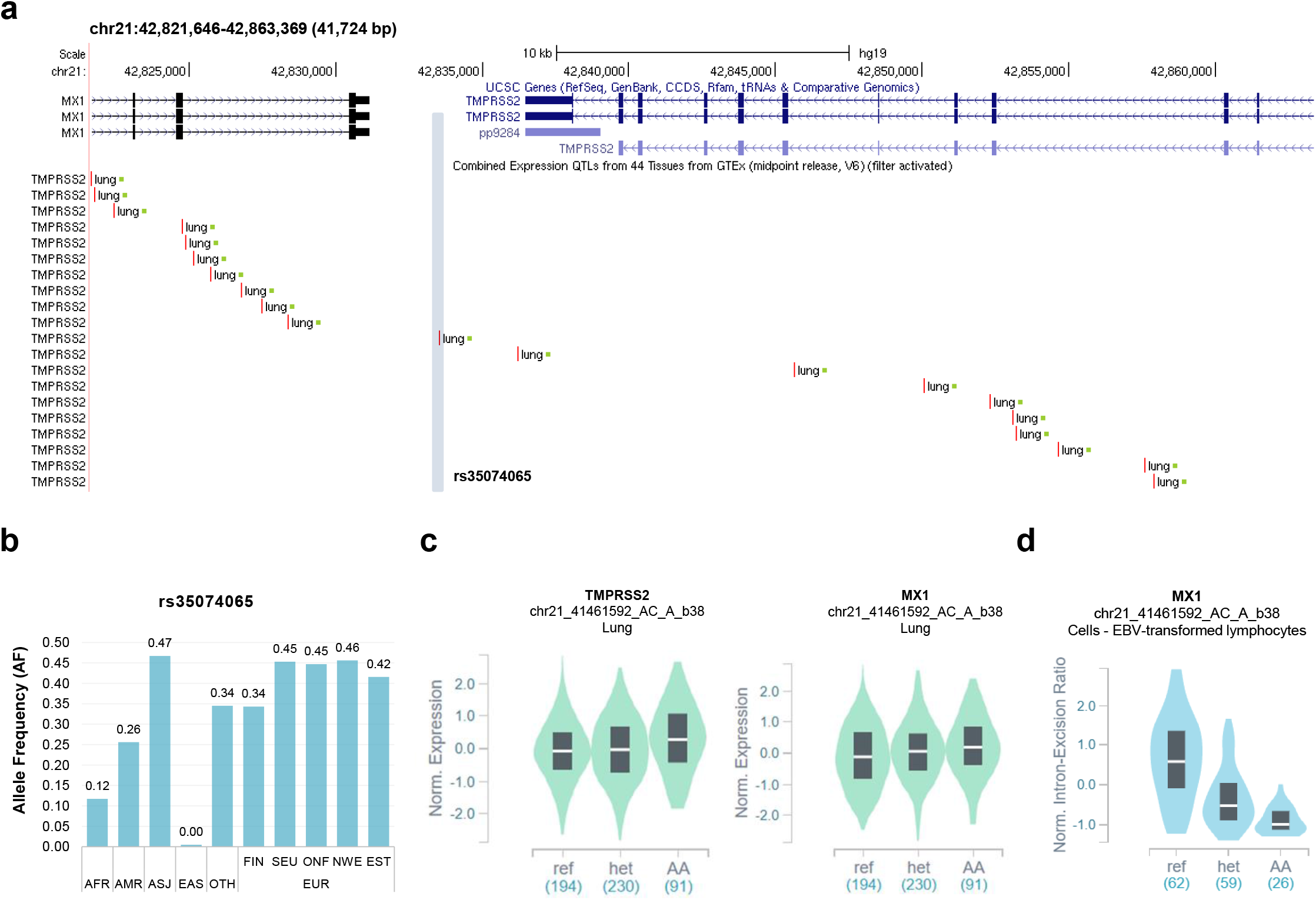
Analysis of the eQTL-lung variants for *TMPRSS2* locus. **a)** Schematics of the genomic region encompassing eQTL lung variants of *TMPRSS2* locus (NES positive ≥ 0.1) by Genome Browser (GRCh37/hg19, https://genome.ucsc.edu/). The most significant eQTL variant is highlighted. **b)** The allele frequencies of del variant rs35074065 were annotated by the gnomAD database (WGS data). AFR, African/African American; AMR, Latino/Admixed American; ASJ, Ashkenazi Jewish; EAS, East Asian; OTH, Other (population not assigned); FIN, Finnish; SEU, Southern European; ONF, Other non-Finnish European; NEW, North-Western European; EST, Estonian. **c)** Violin plot showing the effect of the eQTL rs35074065 variant on *TMPRSS2* and *MX1* expression (*TMPRSS2*: p value = 3.9e-11; NES = 0.13; *MX1:* p value = 0.000010; NES = 0.20). **d)** Violin plot showing the effect of the sQTL rs35074065 variant on MX1 splicing isoform expression (p value = 1.6e-13; NES = –0.83).

## Discussion

Targeting TMPRSS2 expression and/or activity could be a promising candidate for potential interventions against COVID-19 given its crucial role in initiating SARS-CoV-2 and other respiratory viral infections.^16^ Understanding how TMPRSS2 protein expression in the lung varies in the population could reveal important insights into differential susceptibility to influenza and coronavirus infections. Immunohistochemical studies, with limited sample size, suggest that the TMPRSS2 protein is more heavily expressed in bronchial epithelial cells than in surfactant-producing type II alveolar cells and alveolar macrophages, and that there is no expression in type I alveolar cells that form the respiratory surface.^17^ A recent single-cell RNA-sequencing study, confirmed that TMPRSS2 is expressed in type 1 d 2 alveolar cells.^18^ Accordingly, our *in-silico* analysis supported the high *TMPRSS2* gene expression in tissues of the respiratory tract, as bronchus, pharyngeal mucosa, and lung. Moreover, it is also considerable to study the genetic variants and the eQTL of this gene as cause of protein expression variability of TMPRSS2. For example, patients who carried single nucleotide polymorphisms associated with higher TMPRSS2 expression (rs2070788 and rs383510) were more susceptible to influenza virus infection A(H7N9) in two separate patient cohorts.^19^ Our data on eQTL variants showed that the EAS population has much lower AFs in the eQTL lung-specific variants associated with higher *TMPRSS2* expression in lung, while the European populations have higher AFs for the same variants. Interestingly, the top eQTL variants were in a genomic region that includes not only *TMPRSS2* gene but also *MX1* gene, which encodes a guanosine triphosphate (GTP)- metabolizing protein that participates in the cellular antiviral response. *MX1* is an interferon (IFN)-α/β-inducible gene that is widely recognized as an influenza susceptibility gene.^20^ Of note, the downregulation of *MX1* has been documented in non-responder patients to interferon-based antiviral therapy of chronic hepatitis C virus infection.^21^ Our data demonstrated that subjects with the eQTL associated with high expression of *TMPRSS2* could also carry the associated eQTL in the *MX1* gene. In particular, we found that the eQTL variant rs35074065 is associated with high expression of *TMPRSS2* but with a low expression of *MX1* splicing isoform. Thus, these subjects could account for a more susceptibility either to viral infection or to a decrease in cellular antiviral response.

Epidemiological studies across diverse countries including China, Italy, and the United States showed that the incidence and severity of diagnosed COVID-19 as well as other TMPRSS2- dependent viral infections such as influenza may be higher in men than women. Interestingly, we observed the *TMPRSS2* is expressed at high levels in male specific tissues: prostate and testis. In these latter tissues we also found a high number of eQTLs for *TMPRSS2* whose VAF varied among the different population with the lowest frequency in EAS individuals. Another possible explanation of gender differences in mortality and morbidity could be the presence of TMPRSS2:ERG fusion protein in prostate cancer as well as the strong regulation of TMPRSS2 by androgens. Remarkably, at the mRNA level, constitutive expression of TMPRSS2 in lung tissue does not appear to differ between men and women.^16^ Accordingly, *TMPRSS2* gene expression data from GTEx database do not highlight any difference between male in female. There is a wide variation among both sexes in terms of mRNA expression levels.^16^ Low levels of androgens present in women may suffice to sustain TMPRSS2 expression. In addition, TMPRSS2 (and tumors with the TMPRSS2:ERG fusion protein) may be responsive to estrogen signaling.^22,23^ It is attractive to speculate that androgen receptor-inhibitory therapies might reduce susceptibility to COVID-19 pulmonary symptoms and mortality.

In summary, we systematically analyzed coding-region variants in *TMPRSS2* and the eQTL variants, which may affect the gene expression, to compare the genomic characteristics of *TMPRSS2* among different populations. Our findings suggested that no direct evidence was identified genetically supporting the existence of resistant variants for coronavirus S-protein priming in different populations. The effects of low-frequency missense pathogenic variants, as well as those of LoF variants for S-protein priming should be further investigated. The data of variant distribution and AFs may contribute to the further investigations of TMPRSS2, including its roles in acute lung injury and lung function. Of note, our findings suggest that the lung-specific eQTL variants may confer different susceptibility or response to SARS-CoV-2 infection from different populations under the similar conditions. In conclusion, to know the genetic variability of *TMPRSS2* gene will be useful for both the prognosis and the treatment of the patients affected by COVID-19.

## Methods

The variants in *TMPRSS2* gene region (chr21:42836478-42903043, 66.566 Kb) were obtained from the gnomAD v2.1.1 database.^24^ To analyze the distribution of eQTLs for *TMPRSS2*, we used the data from Genotype Tissue Expression (GTEx) database (https://www.gtexportal.org/home/datasets). Annotation of *TMPRSS2* variants and eQTLs was performed with ANNOVAR by using the pathogenicity prediction tools described in Supplementary Table S1 and the allele frequencies of human populations reported in Supplementary Table S5. The reference transcript for *TMPRSS2* annotation was NM_001135099 (ENST00000398585). Genomic coordinates were based on the GRCh37/hg19 build. The classification of non-synonymous variants was performed using the following predictor tools: M-CAP (score >0.025), MutationTester (A-D, disease-causing), CADD v1.3 (Phred score >15) for the pathogenic variants. VEST3 (score <0.5), REVEL (score <0.5), RadialSVM (T, tolerated) were used for the benign variants.^15^ Variants with conflicting interpretation were excluded from further analysis.

## Acknowledgments

The authors thank their colleagues Francesco Manna, Barbara Eleni Rosato, Annalaura Montella e Roberta Marra for continuing their lab work with the same dedication as ever during this troubled period of coronavirus disease COVID-19 pandemic.

## Author Contributions

IA, RR and MC designed and conducted the study, and prepared the manuscript; MC, VAL and RR analysed the data; AI provided critical review of the manuscript.

All the authors read and approved the final manuscript.

