## Supplementary material_Supplementary Figure S1 and Supplementary Figure S2 for "Genetic analysis of the novel SARS-CoV-2 host receptor *TMPRSS2* in different populations"

**Article title:**

**Table of contents:**

- Figure S1
- Figure S2
- Table S1
- Table S2
- Table S3
- Table S4
- Table S5

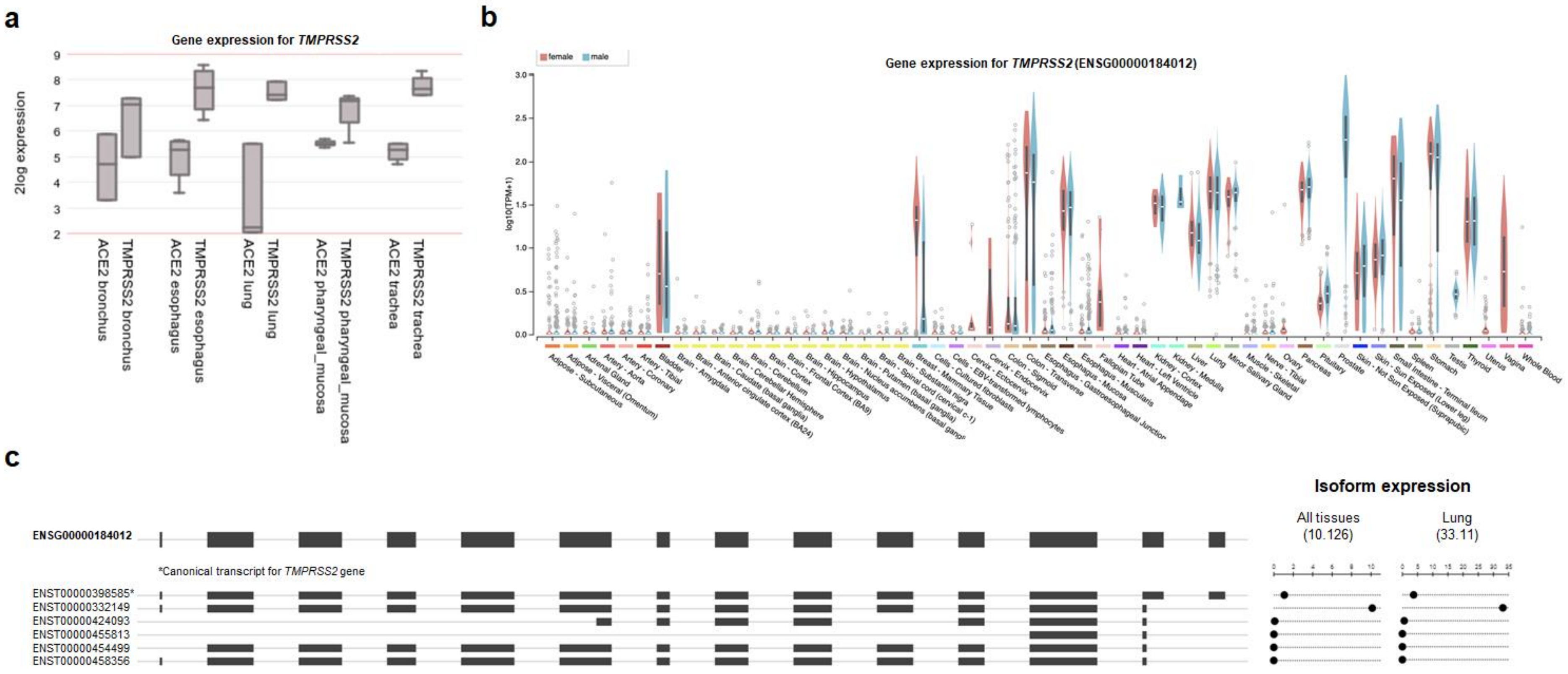


**Figure S1. Gene expression analysis on *TMPRSS2* gene.**

**a)** Expression analysis of the *TMPRSS2* and *ACE* genes by R2, a web-based genomic analysis and visualization tool (R2: Genomics Analysis and Visualization Platform, http://r2.amc.nl). Box plot of each gene expression obtained by the dataset from normal and tumor tissues. Data are shown as 2log expression.

**b)** Gene expression profile of *TMPRSS2* gene in different tissues and stratified according to the gender, as obtained from GTEx database (https://www.gtexportal.org/home).

**c)** Schematics of *TMPRSS2* gene and transcripts as obtained from gnomAD database (https://gnomad.broadinstitute.org/).

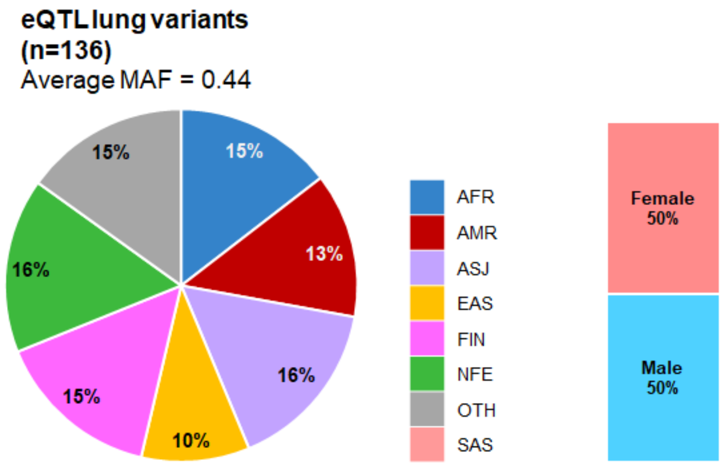


**Figure S2. The allele frequency distribution of eQTL-lung variants of *TMPRSS2* in different populations.** The colors indicate different populations as shown in the color code legend. AFR, African/African American; AMR, Latino/Admixed American; ASJ, Ashkenazi Jewish; EAS, East Asian; FIN, Finnish; NFE, Non-Finnish European; SAS, South Asian; OTH, Other (population not assigned). The histogram on the right shows the AF distribution in overall population stratified according to the gender.
